# Antibody evasion and receptor binding of SARS-CoV-2 LP.8.1.1, NB.1.8.1, XFG, and related subvariants

**DOI:** 10.1101/2025.07.18.662329

**Authors:** Ian A. Mellis, Madeline Wu, Hsiang Hong, Chih-Chen Tzang, Anthony Bowen, Qian Wang, Carmen Gherasim, Virginia M. Pierce, Jayesh G. Shah, Lawrence J. Purpura, Michael T. Yin, Aubree Gordon, Yicheng Guo, David D. Ho

**Author notes:** Correspondence (A.G.), (Y.G.), (D.D.H.). These authors contributed equally (Ian A. Mellis, Madeline Wu, Hsiang Hong, Chih-Chen Tzang, Anthony Bowen).

## Abstract

SARS-CoV-2 continues to evolve, causing repeated waves of infections around the world. It is critical to understand the features of the virus that explain its growth advantages. Recently, the SARS-CoV-2 Omicron JN.1 subvariants KP.3.1.1 and XEC were outcompeted by later JN.1 progenies, most prominently LP.8.1 and LP.8.1.1. Other recent JN.1 subvariants, such as LF.7.2.1, which became prevalent in Asia, and MC.10.1, have also been under monitoring. Subsequently, NB.1.8.1 and XFG subvariants began increasing in prevalence, as well. We found that serum neutralizing antibody titers against LP.8.1, LP.8.1.1, LF.7, LF.7.2.1, MC.10.1 were similar to XEC in a cohort of 20 KP.2-based monovalent mRNA vaccine (KP.2 MV) recipients and in a cohort 20 adults who did not receive KP.2 MV. NB.1.8.1 and XFG were more evasive of serum neutralization than LP.8.1.1. We then characterized subvariant susceptibility to monoclonal antibody (mAb) neutralization using a panel of 12 mAbs spanning several epitopes on the SARS-CoV-2 spike, and found that LP.8.1 and XFG, MC.10.1 and NB.1.8.1, and LF.7.2.1 evade different classes of mAbs relative to earlier JN.1 subvariants, even if the tested polyclonal serum neutralizing antibody titers were not different overall. Next, we found that the receptor-binding affinity of LP.8.1 to ACE2 was the highest among the tested viruses, while that of LF.7.2.1 was lowest. Therefore, unlike most prior SARS-CoV-2 sublineage evolutionary trajectories, receptor-binding affinity, possibly reflecting enhanced transmissibility–and not increased antibody evasion–better explained the rise of LP.8.1, while the expansion of NB.1.8.1 and XFG again appear correlated with their enhanced antibody evasion.

## Introduction

The severe acute respiratory syndrome coronavirus 2 (SARS-CoV-2) Omicron JN.1 subvariants KP.3.1.1 and XEC were dominant around the world until early 2025. However, the virus has continued to evolve rapidly ^1,2^. By March 2025, the JN.1 progeny LP.8.1 and LP.8.1.1 largely replaced XEC in Europe and North America. In parallel, variants such as LF.7 and LF.7.2.1 have increased in frequency across Asia, while MC.10.1 showed a rapid rise in prevalence over a short period, as well. As of June 2025, the NB.1.8.1 and XFG variants have demonstrated a marked growth advantage compared to these aforementioned JN.1 sublineages on a global scale.

LP.8.1 is descended from KP.1, carries five additional spike mutations beyond KP.3.1.1: F186L and R190S in the N terminal domain (NTD), R346T and H445R in the receptor binding domain (RBD), and K1086R in the S2 region. LP.8.1.1 possesses an additional K679R mutation near the furin cleavage site of LP.8.1.1. LF.7 contains seven additional spike mutations beyond JN.1, including four mutations in NTD (T22N, S31P, K182R, and R190S), and three mutations in RBD (R346T, K444R, and F456L). Its descendant, LF.7.2.1, which harbors an additional A475V mutation, has rapidly outcompeted LF.7 in Asia. MC.10.1 was under monitoring because it includes an A435S mutation in the RBD, in addition to the other spike mutations found in KP.3.1.1. NB.1.8.1, a descendant of the recombinant subvariant XDV, subsequently began to rise around the world and carries seven additional mutations beyond JN.1, including T22N, F59S, and G184S in the NTD and A435S, F456L, K478I, and Q493E in the RBD. In addition, XFG, a recombinant subvariant of LF.7 and LP.8.1.2 currently under monitoring and expanding particularly quickly in Europe, bears 4 mutations beyond LF.7: H445R, N487D, and Q493E in the RBD and T572I in SD1. Another recombinant lineage, XFC – derived from LF.7 and LP.8.1.1—retains the NTD mutations from LF.7 and shares other spike mutations with LP.8.1 (Fig. 1A,B).

**Figure 1:**
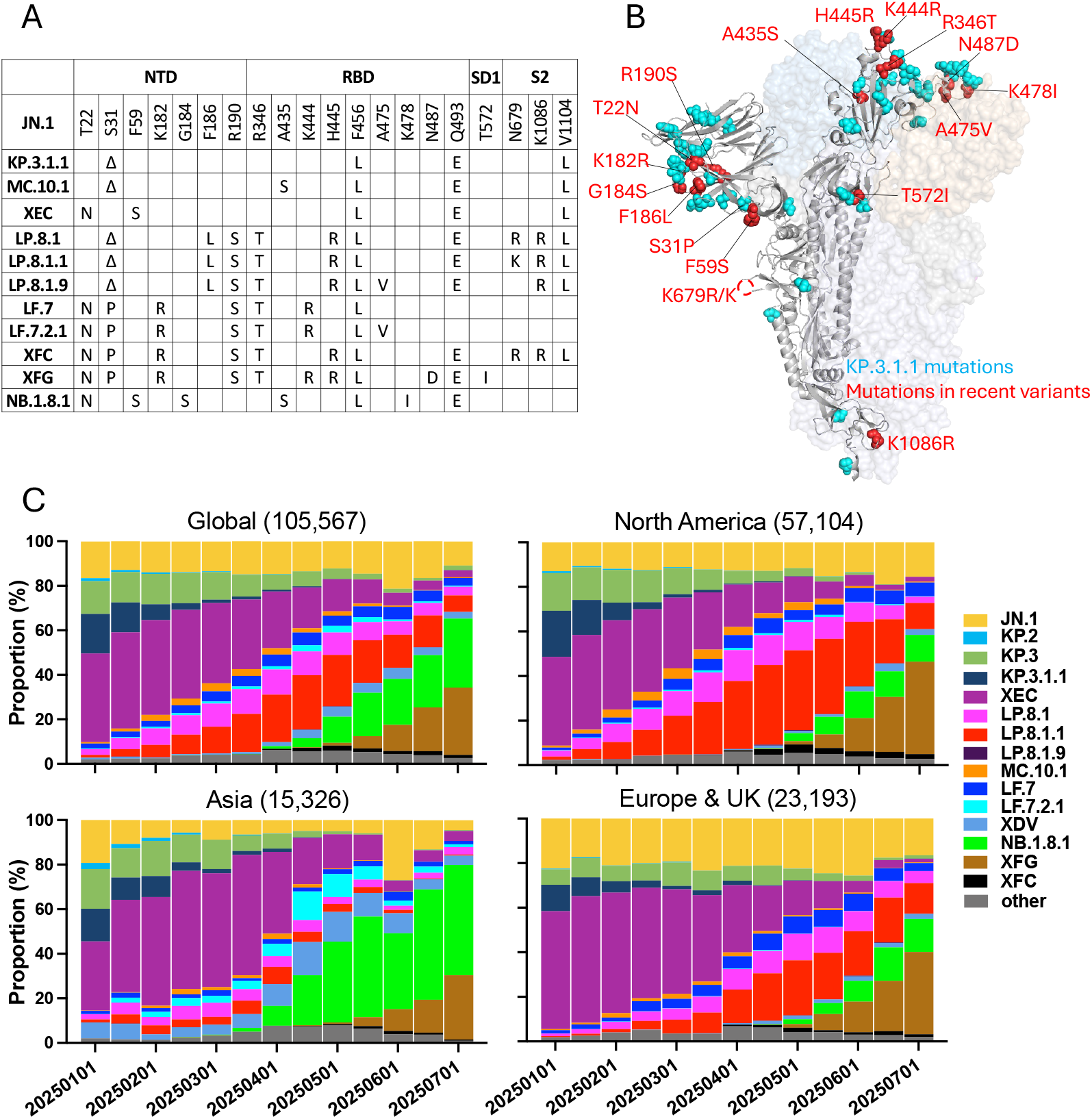
Mutations and frequencies of JN.1 subvariants dominant in Spring 2025. A: Spike mutations present in selected JN.1 subvariants. B: Structural diagram of SARS-CoV-2 spike protein. Spike mutations present in KP.3.1.1 are highlighted in cyan. Spike mutations present in the other JN.1 subvariants studied herein are highlighted in red. C: Relative frequencies of SARS-CoV-2 variants from 01/01/2025 to 07/01/2025 in the indicated regions. Data from GISAID.

Since May 2025, NB.1.8.1 and XFG have expanded globally, emerging from distinct recombinant sublineages of JN.1 (Fig. S1) and demonstrating a clear growth advantage over the previously dominant LP.8.1.1 (Fig. 1C). NB.1.8.1 has rapidly risen to represent over 49% cases in Asia, while XFG accounts for over 36% of new infections in Europe and the UK. XFG is trending to outcompete NB.1.8.1 in North America, where it now accounts for over 40% of cases, compared to only 11% for the latter variant (Fig. 1C).

Given the rapid emergence of these variants, it is critical to characterize the features driving their dominance, including their antigenicity and receptor binding affinity, to inform public health interventions and considerations around vaccine design and to update our understanding of the phenotypic changes driving SARS-CoV-2’s recent evolution. Here, for a panel of pseudoviruses bearing spike proteins of 11 JN.1 subvariants, we compare the viruses’ susceptibility to serum antibody neutralization in two clinical cohorts representative of the population around the time that LP.8.1 rose to dominance in North Americas (with and without KP.2 monovalent vaccine booster). Next, we use neutralization by a panel of monoclonal antibodies (mAbs) to dissect specific epitopes that are altered in dominant viruses. To assess receptor binding affinity, we perform soluble ACE2 inhibition studies. Lastly, we synthesize our results using computational structural modeling of spike mutations that may interfere with antibody or receptor binding.

## Results

### LP.8.1.1, LF.7.2.1, and MC.10.1 are antigenically similar to KP.3.1.1 and XEC

We first asked whether the dominance of LP.8.1.1 and rise of LF.7.2.1 is associated with increased serum neutralizing antibody evasion, as has been the case for many prior dominant SARS-CoV-2 variants^1,3–6^. We collected serum samples from 40 adult participants in the US, including 20 who had received a KP.2-based monovalent mRNA vaccine booster dose approximately 1 month prior (“KP.2 MV” cohort) ^2^, and 20 who had not received a KP.2 booster, during the same time period (“No KP.2 MV” cohort). Participants who did not receive the booster are most representative of the US population, where 2024-25 booster dose uptake was low, at 23.5% of adults by April 26, 2025 ^7^. Within the No KP.2 MV cohort, we included 10 participants who had a reported SARS-CoV-2 infection 1-5 months prior to collection and 10 participants who did not have a known SARS-CoV-2 infection in the preceding 5 months (Tables S1, S2). The average age of participants was 44.5 years and was similar across the two cohorts. Both cohorts had more women than men, with 72.5% of participants being women. We performed pseudovirus neutralization assays in parallel for all participants against a panel of 11 pseudoviruses, bearing the spike proteins of JN.1, KP.2, KP.3, KP.3.1.1, XEC, LP.8.1, LP.8.1.1, LP.8.1.9, MC.10.1, LF.7, or LF.7.2.1. The LP.8.1 descendant LP.8.1.9 has not emerged as dominant, but it does bear the A475V spike mutation found in LF.7.2.1.

In the tested cohorts, we found that there were no dramatic differences in geometric mean serum neutralizing antibody titers (GMT) between XEC and later subvariants, including LP.8.1.1 and LF.7.2.1 (GMT ranges 486-774 in the KP.2 MV cohort and 187-312 in the No KP.2 MV cohort; Fig. 2A,B). In some cases there were small absolute differences that rose to statistical significance, such as the GMT against LP.8.1.1, paradoxically, being higher than against LP.8.1 or XEC (1.2X or 1.4X relative to LP.8.1 in the KP.2 MV cohort). Titers against LF.7.2.1 were the lower end of the range in both cohorts, but were not significantly different from LP.8.1. These findings suggest that the dominance of LP.8.1.1 and expansion of LF.7.2.1 are not associated with dramatic increases in serum antibody evasion in the US. To assess the antigenicity of the tested variants, we generated antigenic maps using the combined set of sera (Fig. 2C). Antigenically, all tested viruses were similar; the tested subvariants were within 1.51 antigenic units of JN.1.

**Figure 2:**
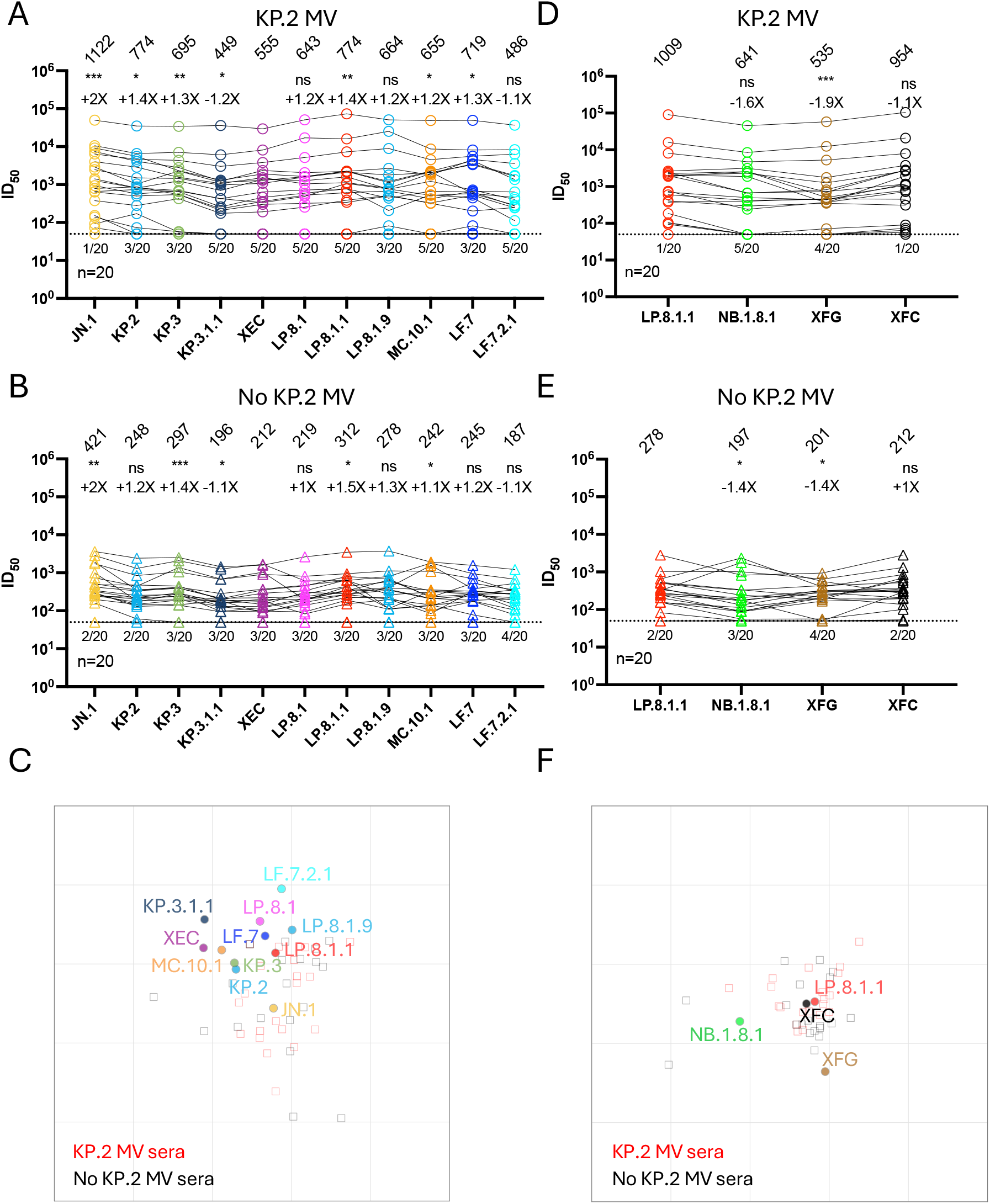
Serum antibody evasion of recent JN.1 subvariants. Serum neutralizing titers (ID_50_) against VSV-based pseudoviruses bearing spike proteins of JN.1, KP.2, KP.3, KP.3.1.1, XEC, LP.8.1, LP.8.1.1, LP.8.1.9, MC.10.1, LF.7, or LF.7.2.1, for samples from 20 recipients of KP.2 MV boosters at ∼1 month post-booster (A; “KP.2 MV”) and 20 US-based adults who chose not to receive a KP.2 MV booster (B; “No KP.2 MV”). The geometric mean ID_50_ titer (GMT) is presented at the top. The fold change in GMT for each virus compared to XEC is also shown immediately above the symbols. Statistical analyses used Wilcoxon matched-pairs signed-rank tests, comparing to XEC. n, sample size; ns, not significant. * p < 0.05, ** p < 0.01, *** p < 0.001, **** p < 0.0001. Numbers under the dotted lines denote numbers of serum samples that were under the limit of detection (ID_50_ < 50). C, Antigenic map generated using all neutralization data from panels A and B. One antigenic unit (AU) represents an approximately 2-fold change in ID_50_ titer. Serum samples and viruses are shown as squares and dots, respectively. Serum neutralizing titers (ID_50_) against VSV-based pseudoviruses bearing spike proteins of LP.8.1.1, NB.1.8.1, XFG, or XFC, for samples from 20 recipients of KP.2 MV boosters at ∼1 month post-booster (D) and 20 US-based adults who chose not to receive a KP.2 MV booster (E). F, Antigenic map generated using all neutralization data from panels D and E.

For a subset of the subvariants, XEC, KP.3.1.1, and LP.8.1, we also asked whether there were differences in the durability of serum neutralizing titers after receiving a KP.2 MV booster dose, to check whether peak titer trends held over time in the KP.2 MV cohort. Therefore, we tested samples at approximately 1 month and 4 months after dosing in a 16-person subset of the KP.2 MV cohort (Figure S2A). We estimated similar estimated half-lives (66-91 days) of serum neutralizing antibody titers against the 3 tested viruses (Figure S2B). Of note, we discussed these KP.2 MV cohort serum neutralizing antibody titer results–but not the No-KP.2-MV cohort results–in a separate preprint focused on vaccine-elicited titers ^2^.

### NB.1.8.1 and XFG are more serum-antibody-evasive than LP.8.1.1

Although the emergence of LP.8.1.1 was not marked by an increase in antibody-evasiveness, we asked whether the newer variants outcompeting it, NB.1.8.1 and XFG, better evaded neutralizing antibodies, in the same cohorts of participants. In the KP.2 MV cohort, geometric mean neutralizing titers against XFG were significantly lower (by 1.9-fold)) than those against LP.8.1.1, while the trend in lower titers against NB.1.8.1 (1.6-fold) did appear significant (Fig. 2D). Of note, however, 5 of the 20 KP.2 MV recipients had titers against NB.1.8.1 below the limit of detection (<50), while only 1 of the 20 was below the limit of detection for LP.8.1.1. We conservatively calculated geometric mean titers by replacing <50 with 50, as described in the Methods. In the No KP.2 MV cohort, titers against both NB.1.8.1 and XFG were significantly lower than against LP.8.1.1 (Fig. 2E). In both cohorts, there was no difference in titers against XFC compared to LP.8.1.1. Antigenically, NB.1.8.1 and XFG were not dramatically distant from LP.8.1.1, at 0.90 and 1.06 antigenic units, respectively (Fig. 2F).

### mAb evasion of recent JN.1 subvariants

Although we did not detect dramatic differences in serum antibody evasion by LP.8.1.1, LF.7.2.1 compared to XEC in the tested cohorts, the mutations in their spikes are in domains that raise concerns that these viruses could evade mAbs in clinical use or development. Furthermore, we asked whether monoclonal antibody neutralization studies could aid in explaining the lower titers seen against the most recent dominant subvariants, NB.1.8.1 and XFG. Therefore, we tested a panel of 12 mAbs with known epitopes, including NTD-SD2-targeting C1717 ^8^, NTD-RBD interface-targeting C68.61 ^9^, RBD Class 1 antibodies BD55-1205, BD55-4637 ^10^, 19-77, 19-77 R71V ^11^, VIR-7229 ^12^; RBD Class 3 antibodies CYFN1006-1 ^13^ and S309 ^14^; and Class 4/1 antibodies 25F9 ^15^, SA55 ^16^, and VYD222 ^17^ in pseudovirus neutralization assays. Relative to XEC, we found that LP.8.1 sublineage viruses, including LP.8.1.1 and LP.8.1.9, and XFG and XFC displayed greater evasion of the tested Class 3 mAbs. LF.7.2.1 more efficiently evaded Class 1 mAbs, critically including VIR-7229, which is presently in clinical trials. LF.7.2.1 also increasingly evaded S309, a Class 3 antibody. MC.10.1 and NB.1.8.1 more efficiently evaded all Class 4/1 antibodies tested than KP.3.1.1 or XEC (Figure 3A). Both tested subvariants bearing the A475V mutation, LP.8.1.9 and LF.7.2.1, increasingly evaded Class 1 antibodies, consistent with escape mutation studies in optimization of 19-77 ^11^.

**Figure 3:**
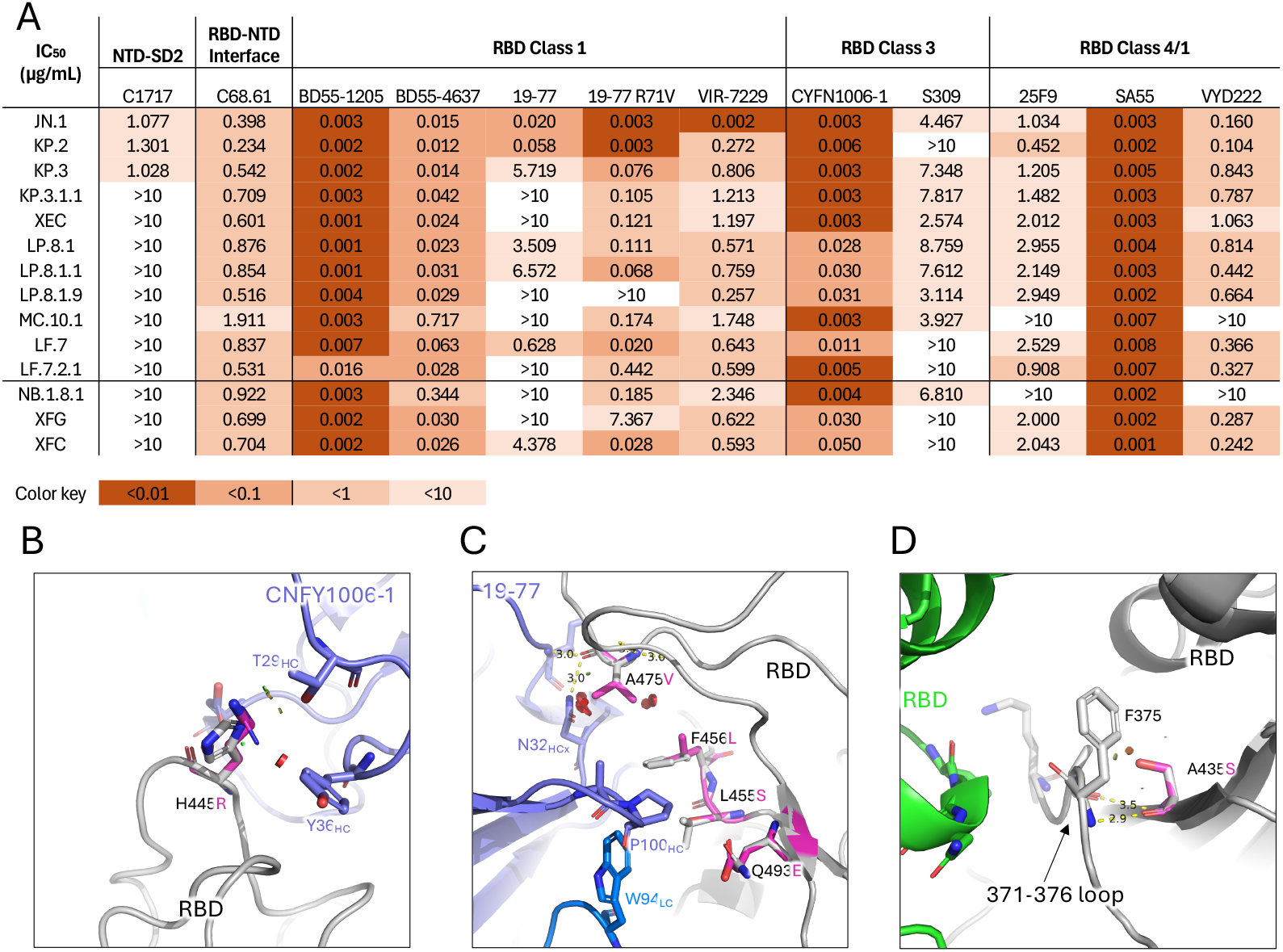
Monoclonal antibody evasion of recent JN.1 subvariants. A: Pseudovirus neutralization IC_50_ values for mAbs against the indicated JN.1 subvariants. B: Modeling of the effects of spike H445R mutation on interaction with CYFN1006-1. C: Modeling of the effects of spike A475V, in the presence of other spike mutations, on interactions with 19-77. D: Modeling of the effects of spike A435S on the spike 371-376 loop.

To explain these results, we performed structural modeling analyses of mAb binding to mutant spikes. We found that the H445R mutation present in LP.8.1, XFG, and XFC introduced minor steric clashes with the CDRH1 of CYFN1006-1 (Fig. 3B), potentially contributing to antibody evasion by LP.8.1 in some individuals by reducing recognition by Class 3 monoclonal antibodies. Additionally, the A475V mutation present in LF.7.2.1 and LP.8.1.9 caused moderate steric hindrance with the 19-77 antibody (Fig. 3C), a representative of VH3-53/66 Class 1 antibodies. Furthermore, the A435S mutation present in MC.10.1 and NB.1.8.1, located near the 371-376 loop of the RBD, introduced a minor steric clash with this loop (Fig. 3D), potentially disrupting its local conformation. As the 371-376 loop plays a key role in modulating RBD structure—as seen originally in Omicron BA.1/BA.2—A435S may alter antigenicity by further inducing conformational changes in the RBD, to make it more often in a “Down” conformation ^18^.

### Receptor binding of recent JN.1 subvariants

Next, we tested whether the spike mutations found in recently dominant variants led to changes in receptor-binding affinity. Therefore, we performed pseudovirus inhibition assays using soluble human ACE2. Relative to XEC, we found that LP.8.1 and LP.8.1.1 had greater receptor-binding affinity (1.77X and 1.59X decreases in IC_50_, respectively; Fig. 4A). However, LF.7.2.1 and MC.10.1 had dramatic impairments in receptor binding, with 4.50X and 2.90X increases in IC_50_ (Fig. 4A). The receptor binding affinity of NB.1.8.1 and XFG, however, was lower than the variant dominant prior to them, LP.8.1.1 (with 1.57X and 2.34X increases in IC_50_).

**Figure 4:**
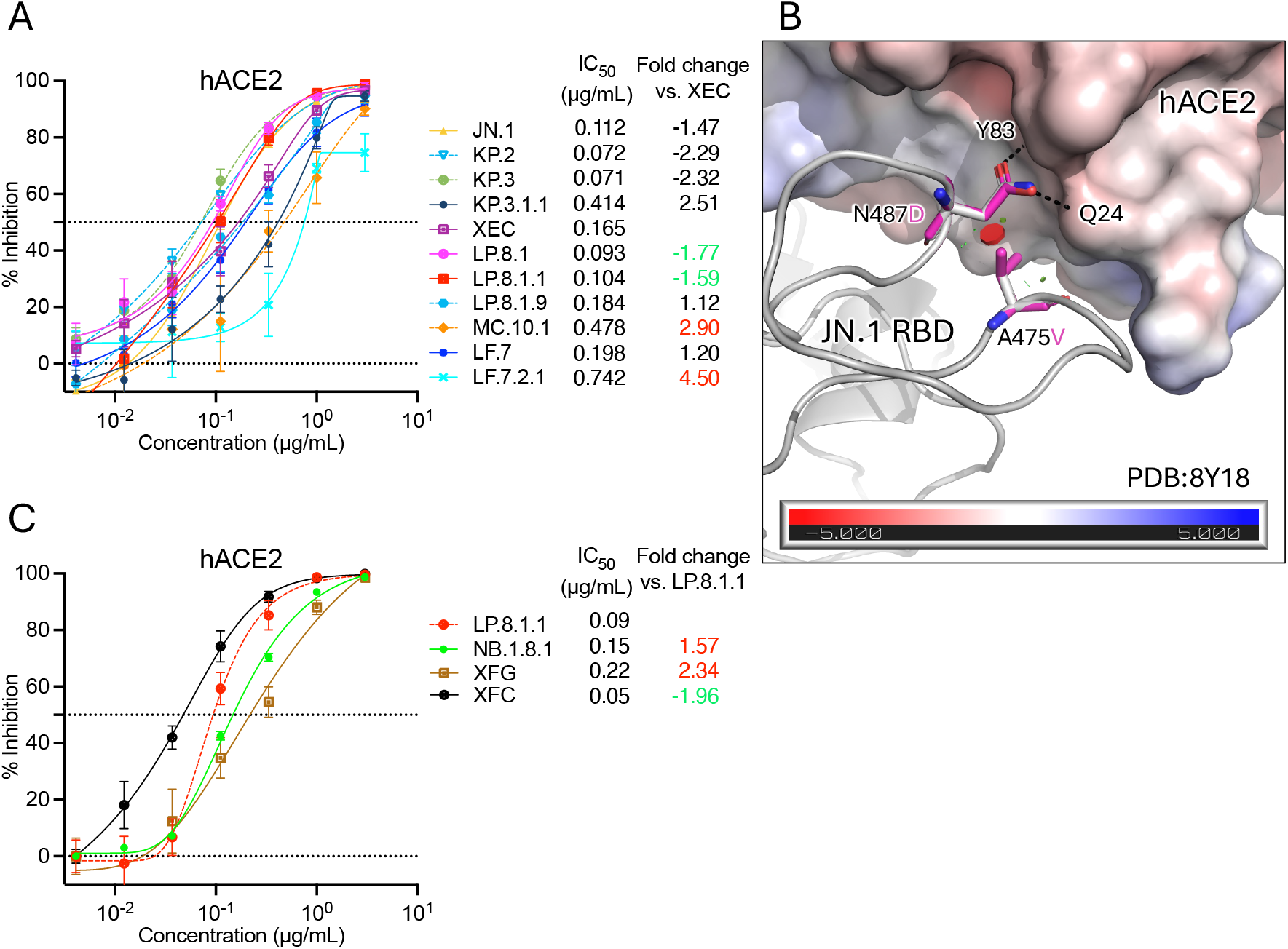
ACE2 binding of recent JN.1 subvariants. A: Sensitivity of 11 JN.1 subvariants to hACE2 inhibition. IC_50_ values are noted, as are fold differences in IC_50_ relative to XEC. B: Modeling of the effects of spike A475V on interactions with spike residue N487, which normally forms hydrogen bonds with hACE2. Electrostatic potential of hACE2 indicated by surface color, according to the scale at bottom, in KT/e. C: Sensitivity of 4 recent JN.1 subvariants to hACE2 inhibition. Data are shown as mean ± standard error of mean (SEM) for three technical replicates.

Structural modeling of the A475V mutation suggests that it may alter the position of RBD residue N487, which normally forms hydrogen bonds with hACE2 (Fig. 4C). In addition, the N487D mutation in XFG introduces a negatively charged residue in place of a neutral one, increasing electrostatic repulsion with the hACE2 receptor, which also has a negatively charged binding interface (Fig. 4C). These structural shifts could explain the reduced ACE2-binding affinity observed in LP.8.1.9, LF.7.2.1, and XFG.

## Discussion

In summary, we found that LP.8.1.1, LF.7.2.1, and MC.10.1 have similar serum neutralizing antibody evasion properties to immediately previously dominant viruses, such as XEC, while the most dominant sublineage of that grouping, LP.8.1, has greater receptor binding affinity.

Subsequent to LP.8.1 and LP.8.1.1, NB.1.8.1 and XFG followed different evolutionary paths by outcompeting LP.8.1.1 likely due to their greater antibody evasion, despite having lower receptor-binding affinity. Therefore, the recent evolution of dominant JN.1 subvariants, since XEC, has been due to multiple factors. At first, dominance was more associated with increasing receptor binding affinity than with evasion of serum neutralizing antibodies, unlike most prior evolutionary trajectories. However, with NB.1.8.1 and XFG, the evolutionary trend returned to prior experience, with antibody evasion becoming most prominent. For example, XEC and KP.3.1.1 were more evasive of serum antibodies than KP.3 and KP.2, and similar trends were observed in the evolution of prior Omicron sublineages, such as XBB and BA.2.75 ^1,3–6^.

LP.8.1 and LP.8.1.1, recent dominant variants globally, and most so in North America, had similar serum antibody evasion properties to prior variants in the tested cohort but bound the receptor ACE2 with greater affinity, consistent with other reports ^19–21^. LF.7.2.1, which was prevalent in Asia but is not expanding as rapidly in North America, also had similar antibody evasion to LP.8.1 and prior variants here, but its receptor binding affinity was much lower than co-circulating strains. This reduction in receptor binding in the absence of dramatic increases in serum antibody evasion among tested North American adults may explain the lack of dominance of LF.7.2.1 in North America. Other groups, using serum from recipients in China and Japan, with different variant exposure and vaccine histories, have observed somewhat lower serum neutralizing antibody titers against LF.7.2.1 in other cohorts, which could explain its different growth advantage in North America vs. Asia ^20,21^. MC.10.1, like LF.7.2.1, had similar serum neutralizing antibody evasion properties, but lower receptor binding affinity, and has been outcompeted by LP.8.1 and other recent variants. Intriguingly, some participants had different trends in titers against the tested variants. For example, in some participants, titers against LP.8.1.1 were higher than against LP.8.1.9, whereas in others, titers against LP.8.1.9 were higher than against LP.8.1.1. Given the variant-specific mAb evasion results above, such results could indicate variability in the anti-spike antibody class composition of individuals’ repertoires. The most concerning recent subvariants, NB.1.8.1 and XFG, are outcompeting LP.8.1.1 around the world, and they most display somewhat greater neutralizing antibody evasion. These results are consistent with one prior report using sera from cohorts in China, however, in a cohort in Japan, titers against NB.1.8.1 were not significantly lower than against LP.8.1 ^22,23^.

Here we have consolidated and presented all recent prominent JN.1 subvariants’ serum and mAb evasion, and estimates of their receptor-binding affinities, for a comprehensive study of the recent evolutionary trajectories of SARS-CoV-2 spike. Overall, these results reveal a temporary shift in correlates of growth advantages from immune escape–primarily mediated by Class 1 and Class 3 monoclonal antibodies evasion–to a more nuanced balance between antibody evasion and receptor-binding affinity. Beginning with the BA.2.86 common ancestor, the virus acquired L455S in JN.1, F456L in KP.2, and Q493E in KP.3—mutations that directly disrupt class 1 antibody binding (Fig. 1A & Fig. 3C). Subsequent sublineages, such as KP.3.1.1 and XEC, accumulated additional changes, including S31del and F59S, which promote an “RBD-down” conformation to further evade class 1 antibodies ^1^. However, this structural shift compromised ACE2 binding affinity, likely imposing a fitness cost (Fig. 4A). To mitigate this, the virus explored multiple evolutionary paths: MC.10.1 and, later, NB.1.8.1 acquired A435S, which reinforced the “RBD-down” state, while LF.7.2.1 and LP.8.1.9 evolved A475V, which directly escaped Class 1 mAbs—though both mutations further reduced receptor affinity. In contrast, LP.8.1 subvariants appear to have restored or enhanced hACE2 binding through R346T, while simultaneously gaining resistance to Class 3 monoclonal antibodies via H445R, contributing to their increased transmissibility and global dominance. These findings underscore the evolutionary plasticity of SARS-CoV-2 and highlight the potential significance of recurrent mutations such as A435S and A475V. Although both are associated with fitness costs, they may become key components of future variants if compensatory mutations emerge to restore or enhance viral fitness. Finally, the recent rise of XFG further supports the notion that SARS-CoV-2 evolution is back to prioritizing immune escape over receptor binding affinity. XFG acquired recurrent mutations R346T and T572I, likely offsetting the detrimental effects of the N487D mutation, which introduces negative charge and electrostatic repulsion at the hACE2 interface ^6^.

### Limitations of this study

The rise of LP.8.1.1 may also be associated with the additional K679R spike mutation near the furin cleavage site, which adds an overlapping cleavage motif, with unknown effects. However, the experiments performed here did not directly address the effects of mutations on S1/S2 cleavage in virion maturation. Furthermore, for all tested variants, we only studied the effects of spike protein mutations. However, mutations in other viral genes could also contribute to growth advantages, though strong such effects have been quite rare in the evolution of SARS-CoV-2.

