## Supplementary Appendix for "Antibody evasion and receptor binding of SARS-CoV-2 LP.8.1.1, NB.1.8.1, XFG, and related subvariants"

**Supplementary Figures and Tables**

**Figure S1: Phylogenetic relationships of JN.1 subvariants.**

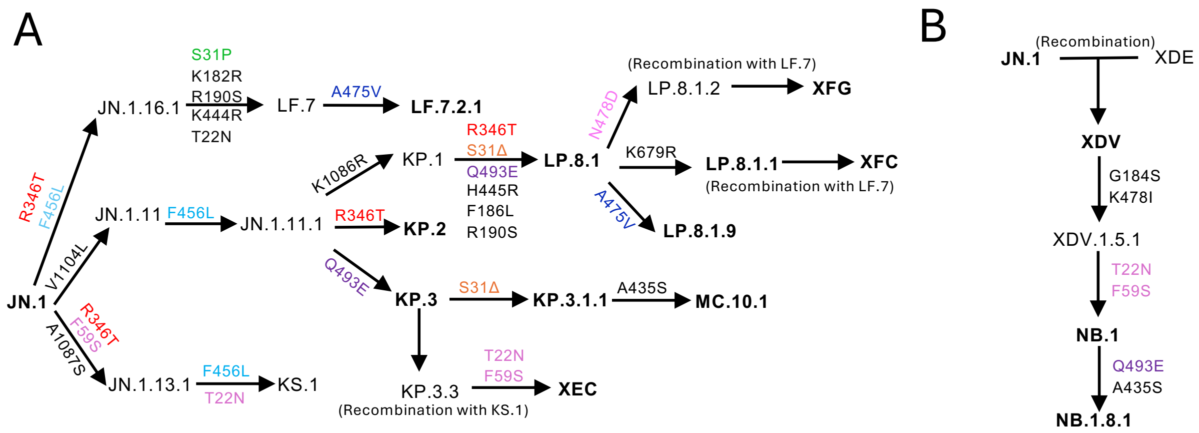

Figure S1: Phylogenetic relationships of JN.1 subvariants. Spike mutations appearing in descendants noted above each arrow. A: Phylogeny of the JN.1 sublineage, excluding NB.1.8.1. B: Recombination event and select ancestors leading to NB.1.8.1.

**Figure S2: Durability of serum neutralizing titers in KP.2 MV recipients.**

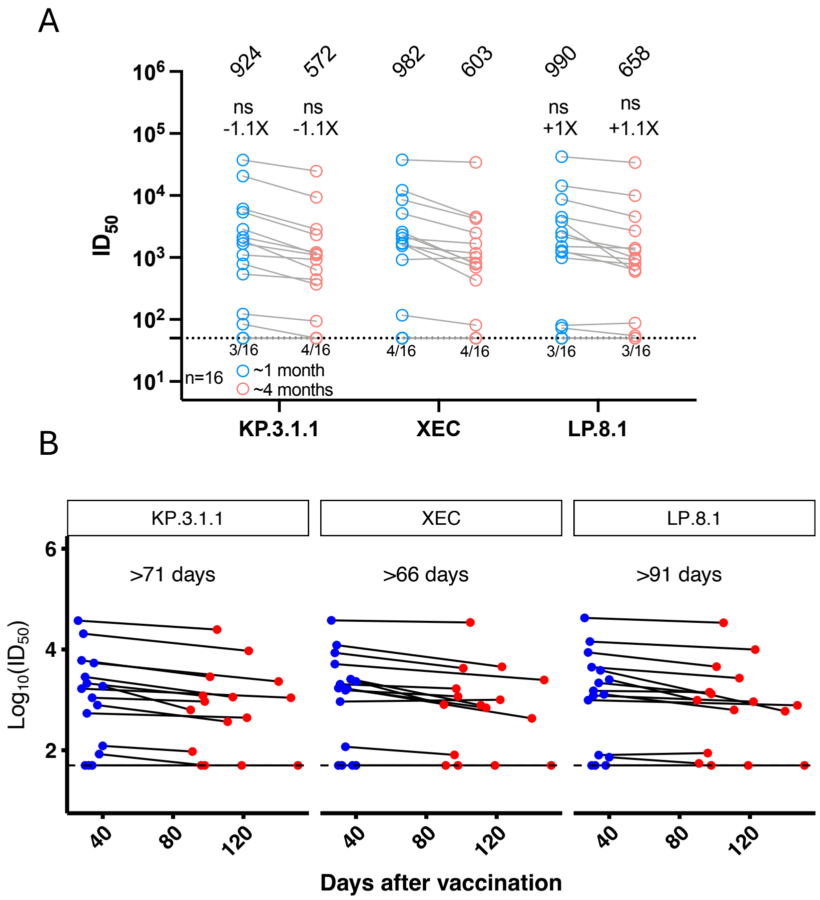

Figure S2: Durability of serum neutralizing antibody titers in KP.2 MV recipients. A: Serum neutralizing titers ($ID_{50}$) against VSV-based pseudoviruses bearing spike proteins of KP.3.1.1, XEC, or LP.8.1, for samples from recipients of KP.2 MV boosters at ~1 month and ~4 months time points post-booster. The geometric mean titer (GMT) is presented at the top. The fold change in GMT for each virus compared to XEC is also shown immediately above the symbols. Statistical analyses used Wilcoxon matched-pairs signed-rank tests, comparing to XEC. n, sample size; ns, not significant. * $p < 0.05$, ** $p < 0.01$, *** $p < 0.001$, **** $p < 0.0001$. Numbers under the dotted lines denote numbers of serum samples that were under the limit of detection (ID50 < 50). B: Paired $ID_{50}$ values per participant. Geometric mean estimated half-life of titers noted for each variant. Half-life was not estimated for participants with unchanged $ID_{50}$ (i.e., both values below LOD), nor for the one participant who had an apparent small increase in titer. Blue = first sampling post-boost, Red = second sampling post-boost.

**Table S1: Summary of clinical cohorts.**

| | | All participants | | KP.2 MV | | No KP.2 MV | |
|---|---|---|---|---|---|---|---|
| | | No. or Mean | % or (range) | No. or Mean | % or (range) | No. or Mean | % or (range) |
| **Total** | | 40 | - | 20 | - | 20 | - |
| **Female** | | 29 | 72.5% | 14 | 70.0% | 15 | 75.0% |
| **Male** | | 11 | 27.5% | 6 | 30.0% | 5 | 25.0% |
| **Age** | | 44.5 | (20, 80) | 42.5 | (21, 80) | 46.5 | (20, 68) |
| | All vaccines | 4.6 | (4, 8) | 5.7 | (4, 8) | 3.5 | (6, 8) |
| | WT | 2.7 | (2, 4) | 3.1 | (2, 4) | 2.3 | (2, 4) |
| | BA.5 BV | 0.7 | (0, 1) | 0.8 | (0, 1) | 0.6 | (0, 1) |
| | XBB.1.5 | 0.7 | (0, 2) | 1.0 | (0, 2) | 0.6 | (0, 2) |
| **No. Vaccines** | KP. 2 MV | 0.5 | (1,1) | 1.0 | (1,1) | - | - |
| **Sera Days Post Most Recent Infection** | | 525.6 | (0, 1685) | 672.8 | (0, 1685) | 404.5 | (0, 1427) |
| **Sera Days Post Most Recent Vaccination** | | 302.5 | (26, 1208) | 33.6 | (26, 50) | 618.8 | (26, 1208) |

**Table S2: Participant demographic, vaccine, and infection details.** Vaccine formulations are denoted as Wild-type (WT), BA.5 Bivalent (BA.5), XBB.1.5 monovalent (XBB.1.5), and KP.2 monovalent (KP.2). Vaccine manufacturers are denoted as Pfizer (P) or Moderna (M).

| ID | Group | Age (Yr) | Sex | Race | No. Vax | No. WT Vax | No. BA.5 Bivalent Vax | No. XBB.1.5 Vax | No. KP.2 MV | Sera Days Post Most Recent Infx | Sera Days Post Most Recent Vaccine | Vaccine History |
|---|---|---|---|---|---|---|---|---|---|---|---|---|
| CUMC 1 | KP.2 MV | 25 | F | As | 6 | 3 | 1 | 1 | 1 | 880 | 31 | WT-P/WT-P/WT-P/BA.5-M/XBB.1.5-M/KP.2-M |
| CUMC 2 | KP.2 MV | 35 | M | As | 4 | 3 | 0 | 0 | 1 | - | 26 | WT-P/WT-P/WT-P/KP.2-M |
| CUMC 3 | KP.2 MV | 21 | M | As | 4 | 3 | 0 | 0 | 1 | 195 | 37 | WT-P/WT-P/WT-P/KP.2-P |
| CUMC 4 | KP.2 MV | 29 | F | As | 4 | 3 | 0 | 0 | 1 | 244 | 28 | WT-P/WT-P/WT-P/KP.2-M |
| CUMC 5 | KP.2 MV | 55 | M | As | 5 | 3 | 1 | 0 | 1 | 451 | 30 | WT-M/WT-M/WT-M/BA.5-P/XBB.1.5-P/KP.2-P |
| CUMC 6 | KP.2 MV | 36 | M | Wh | 6 | 3 | 1 | 1 | 1 | 796 | 30 | WT-P/WT-P/WT-P/WT-P/BA.5-P/XBB.1.5-P/KP.2-M |
| CUMC 8 | KP.2 MV | 23 | F | As | 4 | 3 | 0 | 0 | 1 | - | 40 | WT-P/WT-P/WT-P/KP.2-M |
| CUMC 9 | KP.2 MV | 42 | M | As | 6 | 3 | 1 | 1 | 1 | 650 | 34 | WT-P/WT-P/BA.5-P/XBB.1.5-P/KP.2-M |
| CUMC 11 | KP.2 MV | 61 | F | Wh | 5 | 3 | 1 | 0 | 1 | 826 | 34 | WT-P/WT-P/WT-P/BA.5-P/KP.2-P |
| UMICH 1 | KP.2 MV | 29 | F | Af | 6 | 3 | 1 | 1 | 1 | 784 | 31 | WT-P/WT-P/WT-P/BA.5-P/XBB.1.5-M/KP.2-P |
| UMICH 2 | KP.2 MV | 24 | F | Wh | 6 | 3 | 1 | 1 | 1 | 658 | 28 | WT-P/WT-P/WT-P/BA.5-P/XBB.1.5-M/KP.2-P |
| UMICH 3 | KP.2 MV | 25 | F | Wh | 6 | 3 | 1 | 1 | 1 | - | 32 | WT-P/WT-P/WT-M/BA-M/XBB.1.5-M/KP.2-P |
| UMICH 4 | KP.2 MV | 80 | F | Wh | 8 | 4 | 1 | 2 | 1 | - | 40 | WT-P/WT-P/WT-P/WT-P/BA.5-P/XBB.1.5-P/XBB.1.5-P/KP.2-P |
| UMICH 5 | KP.2 MV | 33 | F | As | 6 | 3 | 1 | 1 | 1 | - | 38 | WT-P/WT-P/WT-P/BA.5-M/XBB.1.5-P/KP.2-P |
| UMICH 6 | KP.2 MV | 55 | F | Wh | 6 | 3 | 1 | 1 | 1 | 653 | 29 | WT-M/WT-M/WT-M/BA.5-M/XBB.1.5-M/KP.2-M |
| UMICH 7 | KP.2 MV | 59 | F | Wh | 7 | 4 | 1 | 1 | 1 | - | 35 | WT-P/WT-P/WT-P/WT-P/BA.5-P/XBB.1.5-M/KP.2-P |
| UMICH 8 | KP.2 MV | 64 | M | Wh | 6 | 3 | 1 | 1 | 1 | 741 | 31 | WT-P/WT-P/WT-P/BA.5-P/XBB.1.5-P/KP.2-P |
| UMICH 9 | KP.2 MV | 66 | F | Wh | 6 | 3 | 1 | 1 | 1 | 1685 | 31 | WT-M/WT-M/WT-P/BA.5-P/XBB.1.5-P/KP.2-P |
| UMICH 10 | KP.2 MV | 27 | F | Wh | 5 | 2 | 1 | 1 | 1 | 476 | 37 | WT-J/WT-M/BA.5-M/XBB.1.5-M/KP.2-M |
| UMICH 11 | KP.2 MV | 61 | F | Wh | 7 | 4 | 1 | 1 | 1 | 380 | 50 | WT-P/WT-P/WT-P/WT-P/BA.5-P/XBB.1.5-P/KP.2-M |
| UMICH 12 | No KP.2 MV | 41 | F | Wh | 2 | 2 | 0 | 0 | 0 | 142 | 1208 | WT-P/WT-P |
| UMICH 13 | No KP.2 MV | 72 | F | Wh | 6 | 3 | 1 | 2 | 0 | 109 | 139 | WT-P/WT-P/WT-P/BA.5-P/XBB.1.5-P/XBB.1.5-P |
| UMICH 14 | No KP.2 MV | 60 | F | Wh | 5 | 3 | 1 | 1 | 0 | 573 | 207 | WT-M/WT-M/WT-M/BA.5-M/XBB.1.5-M/XBB.1.5-M |
| UMICH 15 | No KP.2 MV | 68 | F | Wh | 5 | 3 | 1 | 1 | 0 | 785 | 298 | WT-M/WT-M/WT-M/BA.5-M/XBB.1.5-M/XBB.1.5-M |
| UMICH 16 | No KP.2 MV | 40 | F | Wh | 0 | 0 | 0 | 0 | 0 | - | - | - |
| UMICH 17 | No KP.2 MV | 36 | M | Wh | 5 | 3 | 1 | 1 | 0 | 1208 | 368 | WT-P/WT-P/WT-P/BA.5-P/XBB.1.5-P |
| UMICH 18 | No KP.2 MV | 23 | M | As | 3 | 3 | 0 | 0 | 0 | 90 | 1033 | WT-P/WT-P/WT-P |
| UMICH 19 | No KP.2 MV | 33 | F | As | 5 | 3 | 1 | 1 | 0 | 271 | 383 | WT-P/WT-P/WT-P/BA.5-P/XBB.1.5-P |
| UMICH 20 | No KP.2 MV | 20 | F | Wh | 3 | 1 | 1 | 1 | 0 | 59 | 364 | WT-P/BA.5-M/XBB.1.5-M |
| UMICH 21 | No KP.2 MV | 61 | M | Wh | 5 | 3 | 1 | 1 | 0 | 109 | 402 | WT-P/WT-P/WT-P/BA.5-P/XBB.1.5-P |
| UMICH 22 | No KP.2 MV | 48 | F | Af | 4 | 3 | 0 | 1 | 0 | 109 | 414 | WT-P/WT-P/WT-P/XBB.1.5-P |
| UMICH 23 | No KP.2 MV | 64 | F | Wh | 2 | 2 | 0 | 0 | 0 | - | 1092 | WT-M/WT-M |
| UMICH 24 | No KP.2 MV | 35 | F | Wh | 3 | 3 | 0 | 0 | 0 | 1427 | 1163 | WT-P/WT-P/WT-P |
| UMICH 25 | No KP.2 MV | 61 | M | Wh | 4 | 3 | 1 | 0 | 0 | 972 | 789 | WT-M/WT-M/WT-M/BA.5-P |
| UMICH 26 | No KP.2 MV | 42 | F | As | 5 | 3 | 1 | 1 | 0 | 155 | 429 | WT-P/WT-P/WT-P/BA.5-P/XBB.1.5-P |
| UMICH 27 | No KP.2 MV | 42 | F | Af | 0 | 0 | 0 | 0 | 0 | 507 | - | - |
| UMICH 28 | No KP.2 MV | 50 | F | Wh | 3 | 3 | 0 | 0 | 0 | 37 | 1164 | WT-P/WT-P/WT-P |
| UMICH 29 | No KP.2 MV | 44 | F | Other | 4 | 2 | 1 | 1 | 0 | 88 | 515 | WT-U/WT-P/BA.5-P/XBB.1.5-M |
| UMICH 30 | No KP.2 MV | 36 | F | Wh | 0 | 0 | 0 | 0 | 0 | Unknown | - | - |
| UMICH 31 | No KP.2 MV | 53 | M | Wh | 5 | 3 | 1 | 1 | 0 | 235 | 552 | WT-P/WT-P/WT-P/BA.5-P/XBB.1.5-P |

### Supplementary Methods

#### Clinical Cohorts

Serum samples were collected through the VIVA study at the University of Michigan and through the "COVID-19 Persistence and Immunology Cohort (C-PIC)" study at Columbia University. Specimens were obtained following participant informed consent, adhering to the protocols approved by the IRBs of University of Michigan Medical School (protocol HUM00232359) and Columbia University (protocol AAAS9722).

In this study, serum samples were collected from individuals who had been administered the KP.2 monovalent vaccine booster (KP.2 MV) and from those who had chosen not to receive a KP.2 MV. In the KP.2 MV cohort, serum was collected at approximately 1 month and, for a subset of the participants, also at approximately 4 months after receiving the booster. In the No KP.2 MV cohort, participants were eligible to participate if they had not had a documented SARS-CoV-2 infection within the last month. 10 of the 20 participants in the No KP.2 MV cohort reported an infection between 5 months and 1 month prior to collection (as early as November 2024), and the other 10 participants in the No KP.2 MV cohort reported no recent infections.

The majority of study subjects were female, 72.5%, with an average age of 44.5 years. Serum samples were collected, on average, 33.6 days and, for those with a second sampling, 112.7 days post KP.2 MV booster. Further demographic details, vaccination status, and serum collection timelines are summarized in Tables S1 and S2. NP ELISAs were performed, as previously described, to check for evidence of unreported infections between the two samples for each participant in the KP.2 MV cohort with 1- and 4-month sample pairs. All serum samples were heat inactivated at 56°C for 30 min before use.

#### Cell lines

Vero-E6 (CRL-1586) cells and HEK293T (CRL-3216) cells were obtained from ATCC and cultured at 37°C with 5% CO2 in Dulbecco's Modified Eagle Medium (DMEM) + 10% fetal bovine serum (FBS) + 1% penicillin-streptomycin. Expi293 (A14527) cells were purchased from Thermo Fisher Scientific and maintained in Expi293 expression medium per the manufacturer's instructions. Vero-E6 cells are derived from African green monkey kidneys. HEK293T cells and Expi293 cells are of human female origin.

#### Plasmid generation

As previously described, antibody sequences for the heavy chain variable (VH) and the light chain variable (VL) domains were synthesized by GenScript and then cloned into the gWiz vector to produce antibody expression plasmids. For the packaging plasmids for pseudoviruses, mutations were made by using the QuikChange II XL and QuikChange Multi site-directed mutagenesis kits (Agilent). All constructs were verified using Sanger sequencing prior to use.

### Protein expression and purification

The gWiz-antibody, pcDNA3-sACE2-WT(732)-IgG1 (Addgene 154104) plasmid was transfected into Expi293 cells using PEI at a ratio of 1:3, and then the supernatants were collected after five days. The antibodies and human ACE2 (hACE2) fused to a Fc tag were purified with Protein A Sepharose (Cytiva) following the manufacturer's instructions. Molecular weight and purity were confirmed by SDS-PAGE protein electrophoresis prior to use.

### Pseudovirus production

VSV-based SARS-CoV-2 pseudoviruses, in which the native VSV glycoprotein was replaced by SARS-CoV-2 spike and its variants, were produced as previously described (3). Briefly, plasmids containing the appropriate spike were transfected into HEK293T cells with PEI. After 24 hours, VSV-G pseudotyped ΔG-luciferase (G*ΔG-luciferase, Kerafast) was added, and then washed with medium three times before being cultured in fresh medium for another 24 hours. Anti-VSVG (anti-I1) antibody was added to deplete non-pseudotyped viruses. Pseudoviruses were then harvested, centrifuged, and then aliquoted and stored at -80°C.

### Pseudovirus neutralization assays with sera, mAbs, or ACE2

Each SARS-CoV-2 pseudovirus was titrated to standardize viral infectious dose before use in neutralization assays. Seven serial dilutions of heat-inactivated sera, monoclonal antibodies (mAbs), of soluble ACE2 were added in 96-well plates, starting at 1:50 dilution for sera, 10 μg/mL for antibodies, and 3 μg/mL for ACE2. For ACE2 inhibition assays, as previously reported, we used soluble chimeric human ACE2, which contains ACE2 residues 1-732 fused to human IgG1 Fc. Next, pseudoviruses were added and incubated at 37 °C for 1 hour. In each plate, wells containing only pseudoviruses were included as controls. $4 \times 10^4$ Vero-E6 cells were then added per well and incubated at 37 °C for 16 hours. Promega Luciferase Assay System (E4550) was used for lysis and luciferase activity measurements on a Tecan Infinite® 200 PRO using i-control™ software v.3.9.1.0, in accordance with the manufacturer's instructions. The serum dilution, mAb concentration, or hACE2 concentration that inhibits 50% of virus entry ($ID_{50}$ or $IC_{50}$) was calculated using five-parameter log-logistic dose-response curve fitting with the drda package (v2.0.5) in R.

### Antigenic cartography

Antigenic distances between sera, JN.1 and JN.1 subvariants were determined by integrating all $ID_{50}$ values of individual serum samples through a published antigenic cartography approach (4). The visualization was generated using Racmacs (v.1.1.4, https://acorg.github.io/Racmacs/) in R version 4.3.2. The optimization step count was set at 2,000 and the minimum column basis parameter set to 'none', the 'mapDistances' function was employed to calculate antigenic distances between each serum sample and variant.

### Quantification and statistical analysis

Neutralization $ID_{50}$ and $IC_{50}$ values were determined by fitting a five-parameter dose-response curve in GraphPad Prism v9.3. Statistical significance of differences in neutralizing titer was evaluated using two-tailed Wilcoxon matched-pairs signed-rank tests in GraphPad Prism v9.3. Significance is presented as following: ns, not significant; *p < 0.05; **p < 0.01; and ***p < 0.001, and ****p < 0.0001.

### Author Contributions

The study was conceptualized by I.A.M., Y.G., and D.D.H. Experiments were conducted and data analyzed by I.A.M., M.W., H.H., C.-C.T., Q.W., and Y.G. Project management was handled by I.A.M. Serum samples were collected by I.A.M., A.B., C.G., V.M.P., J.G.S., L.J.P., M.T.Y., A.G., and their colleagues. The results were analyzed, and the manuscript was written by I.A.M., M.W., Y.G., and D.D.H. All contributing authors have reviewed and endorsed the manuscript.

### Declaration of Interests

D.D.H. co-founded TaiMed Biologics and RenBio, and he serves as a consultant for WuXi Biologics and Brii Biosciences and is a board director at Vicarious Surgical. A.G. served as a member of the scientific advisory board for Janssen Pharmaceuticals and has consulted and serves on a scientific advisory board for Sanofi Pasteur. The remaining authors declare no conflicts of interest.

### Acknowledgements

This study was supported by funding from the NIH SARS-CoV-2 Assessment of Viral Evolution (SAVE) Program (subcontract no. 0258-A700-4609 under federal contract no. 75N93021C00014 to D.D.H. and (subcontract GR0010139-PO024016 under federal contract no. 75N93021C00016) to A.G. and the Gates Foundation (project INV019355) to D.D.H., internal startup funding UR014016 from Columbia University to Y.G. K08 AI180347 to A.B., K23 AI171263 to L.J.P., K24 AI155230 to M.T.Y. We thank all who contributed their data to the Global Initiative on Sharing All Influenza Data (GISAID).

We thank Amanda Castillo, Meredith McNairy and Antonia Sturiza for conducting the C-PIC study (Columbia), and to Zijin Chu, Theresa Kowalski-Dobson, Anna Buswinka, Gabe Simjanovski, Joseph Wendzinski, Mayurika Patel, Kathleen Lindsey, and Dawson Davis of the VIVA study team for conducting the VIVA study (at University of Michigan).
